# A minimal role for synonymous variation in human disease

**DOI:** 10.1101/2022.07.13.499964

**Authors:** Ryan S. Dhindsa, Quanli Wang, Dimitrios Vitsios, Oliver S. Burren, Fengyuan Hu, James E. DiCarlo, Leonid Kruglyak, Daniel G. MacArthur, Matthew E. Hurles, Slavé Petrovski

## Abstract

Synonymous mutations change the DNA sequence of a gene without affecting the amino acid sequence of the encoded protein. Although emerging evidence suggests that synonymous mutations can impact RNA splicing, translational efficiency, and mRNA stability^1^, studies in human genetics, mutagenesis screens, and other experiments and evolutionary analyses have repeatedly shown that most synonymous variants are neutral or only weakly deleterious, with some notable exceptions. In their recent article, Shen et al. claim to have disproved these well-established findings. They perform mutagenesis experiments in yeast and conclude that synonymous mutations frequently reduce fitness to the same extent as nonsynonymous mutations^2^. Based on their findings, the authors state that their results “imply that synonymous mutations are nearly as important as nonsynonymous mutations in causing disease.” An accompanying News and Views argues that “revising our expectations about synonymous mutations should expand our view of the genetic underpinnings of human health”^3^. Considering potential technical concerns with these experiments^4^ and a large, coherent body of knowledge establishing the predominant neutrality of synonymous variants, we caution against interpreting this study in the context of human disease.

## Main Text

Purifying selection typically removes deleterious variants from a population before they can reach a high frequency. If most synonymous variants have deleterious fitness effects similar to those of nonsynonymous variants, as Shen et al. claim, then these two classes of variants should appear at similar allele frequencies. However, large population genetics studies across multiple species have repeatedly demonstrated that this is not the case. In yeast, nonsynonymous mutations display a stronger bias toward rare alleles than do synonymous variants across multiple strains^5,6^ and are more likely to contribute to phenotypic variation^7^. Likewise, analyses of allele frequency spectra in Drosophila and in E. coli have suggested that most synonymous sites are subject to very weak, if any, selection^8,9^.

In humans, synonymous variants are expected to be under even less selection as a consequence of a small effective population size. Indeed, in our previous analysis of ∼60,000 whole genomes in the TopMED database^10^, the site frequency spectrum (SFS) of synonymous variants was nearly identical to the SFS of intronic variants^11^. On the other hand, missense variants and loss-of-function variants appear at much lower allele frequencies than synonymous variants. This observation holds true even when accounting for synonymous variants expected to affect mRNA levels via changes in codon optimality. In the gnomAD dataset of ∼123,000 exomes^12^, we found that although codon optimality-reducing synonymous variants appear to be under purifying selection, they are under significantly weaker selection than are missense and protein-truncating variants^11^.

Motivated by the claims of Shen et al., we revisited this analysis in a much larger sample of 454,668 human exomes available in the UK Biobank to show that synonymous variants explain a demonstrably smaller proportion of the genetic architecture of human traits than do nonsynonymous variants, consistent with the prevailing view in the field. We used the codon stability coefficient (CSC), which measures the effect of synonymous codons on mRNA half-life in human cell lines, to determine the codon optimality of fourfold degenerate synonymous codons^13^. Consistent with prior findings, we found that protein-truncating variants (PTVs) and missense variants both appear at lower frequencies than codon-optimality-reducing synonymous variants (P<1×10^−300^ for both comparisons; **Figure 1**). Synonymous variants that reduce codon optimality did appear at lower frequencies in human populations than synonymous variants that do not change optimality (i.e., “neutral” variants, Wilcoxon P=6.2×10^−27^). However, their allele frequency distribution remained strikingly different from nonsynonymous variants (**Figure 1**). These data firmly demonstrate that synonymous variants are under significantly weaker constraint than nonsynonymous variants and thus more likely to be neutral. In support of this, multiplexed assays of variant effect (MAVEs) have shown that nonsynonymous variants are on average far more deleterious than synonymous variants in both humans and yeast^14^ (**Extended Data Figure 1**).

**Figure 1.**
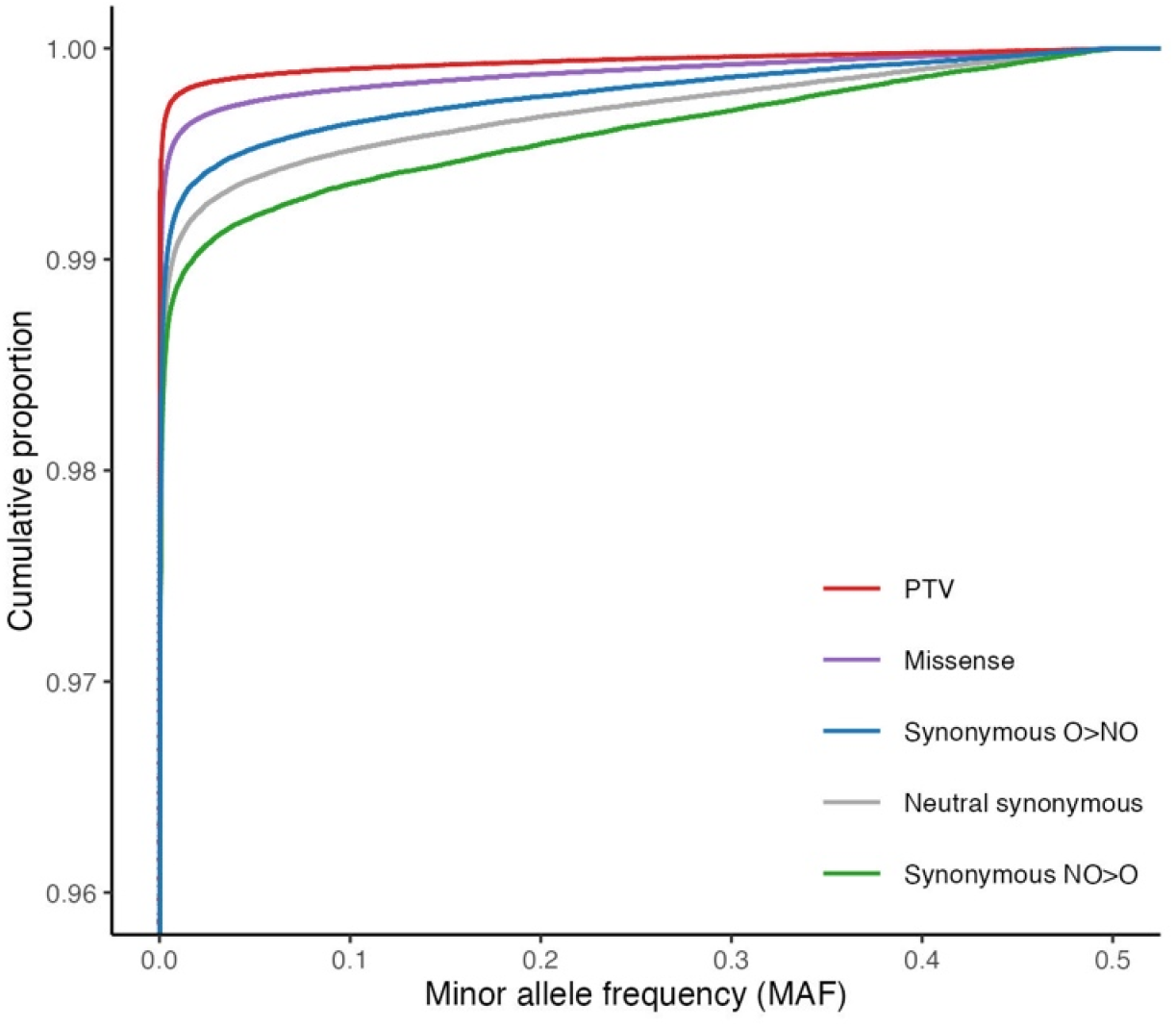
Allele frequency spectrum of synonymous and nonsynonymous variants in ∼450,000 UK Biobank participants. UK Biobank allele frequencies of protein-truncating variants (PTVs; n=913,315), missense variants (n=7,215,418), synonymous optimal-to-nonoptimal variants (O>NO; n=750,732), neutral synonymous variants (i.e., those that do not change optimality; n=843,191), and nonoptimal-to-optimal synonymous variants (n=338,387).

Moreover, genetic association studies and clinical sequencing studies that link genetic variation to clinical disease have consistently demonstrated that nonsynonymous variants have a greater impact on the genetic architecture of human diseases than synonymous variants. For example, large trio-based sequencing studies assessing the contributions of *de novo* mutations in epileptic encephalopathies, autism spectrum disorder, and developmental disorders have identified hundreds of genes with a significant enrichment of *de novo* nonsynonymous variants, but none with a significant enrichment of synonymous variants^15–18^. Similarly, in the setting of cancer, driver mutations are more significantly enriched for missense variants and PTVs than synonymous variants^19^.

To further assess the relative contribution of synonymous and nonsynonymous variants across a wider range of common human phenotypes, we leveraged our published phenome-wide gene-based collapsing analysis performed on 394,694 UK Biobank participants (https://azphewas.com)^20^. For this analysis, we compared the number of significant gene-level associations between 18,762 genes and 4,911 clinical phenotypes using different qualifying variant models. Here, we compare the number of significant (P<2×10^−9^) gene-phenotype associations that emerged from the synonymous variant model, the “raredmg” missense model that includes putatively damaging missense variants, and the PTV model (see Methods; **Fig. 2a**). Across 18,762 genes, only two genes were significantly associated with at least one phenotype in the synonymous model, compared to 32 genes (16-fold enrichment) in the damaging missense model, and 55 genes (28-fold enrichment) in the PTV model (**Fig. 2b**). These data provide yet another empirical validation of the more substantial role that nonsynonymous variation plays in human disease.

**Figure 2.**
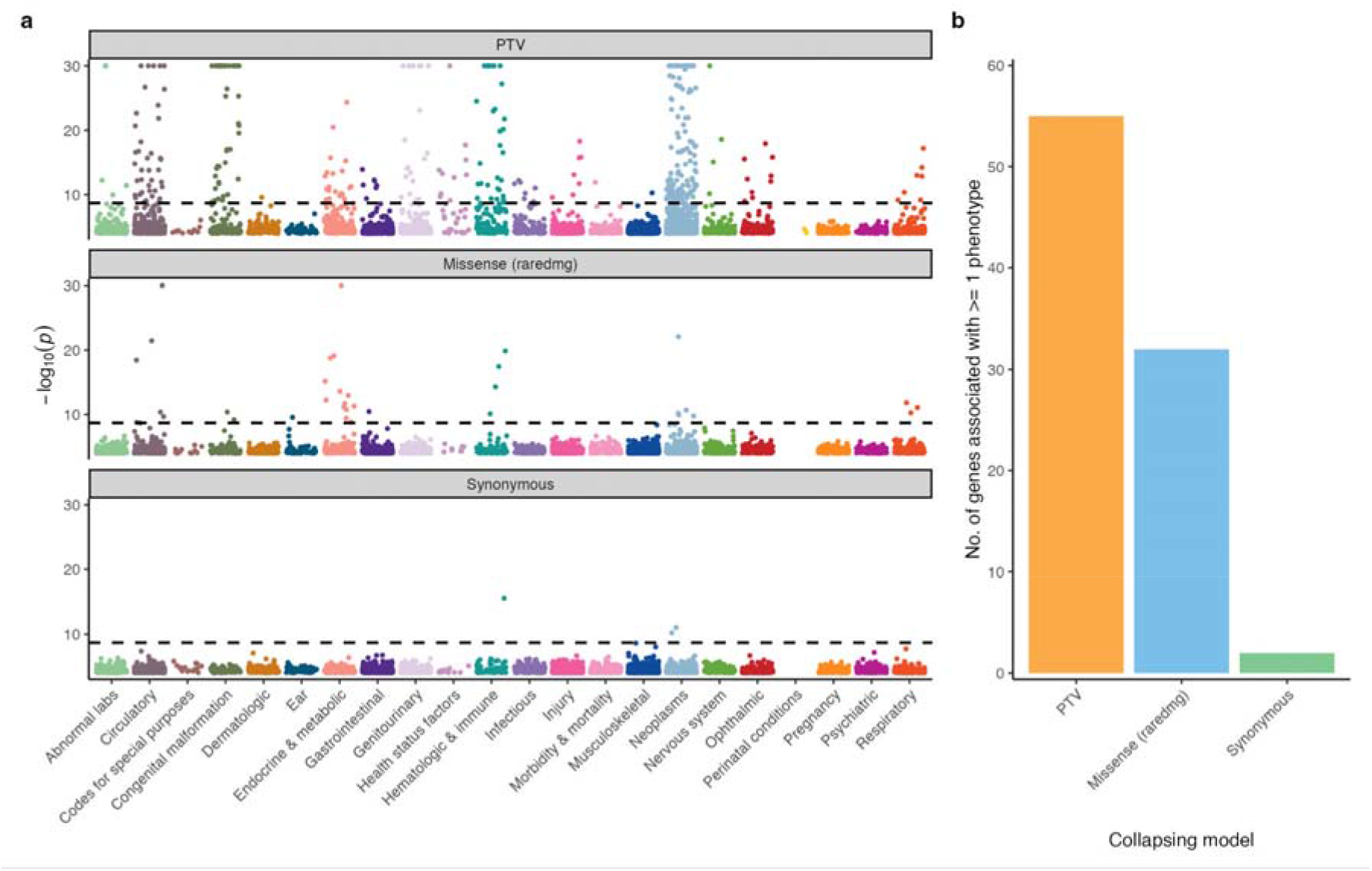
UK Biobank phenome-wide collapsing analysis. **(a)** Gene-based collapsing analysis across 4,911 UK Biobank phenotypes, classified by ICD10-based chapters. The PTV model includes PTVs with MAF<0.1%. The missense (raredmg) model includes missense variants with a MAF<0.005% and REVEL>0.25^20^. The synonymous model includes synonymous variants with MAF<0.005%. **(b)** The number of genes per collapsing model that were significantly (p<2×10-9) associated with at least one phenotype (PTV n=55, Missense n=32, synonymous n=2).

Collectively, evidence from population genetic studies of purifying selection, disease-causing mutations in clinical cohorts, and association studies in population biobanks show that synonymous variants are generally more likely to be neutral than nonsynonymous variants. The higher allele frequencies of synonymous variants compared to nonsynonymous variants across both humans and yeast strongly suggest that the contradictory findings in Shen et al. are not attributable to differences in species. Importantly, our results do not imply that all synonymous variants are strictly neutral, and there are some examples of synonymous variants associated with human traits^21,22^; rather, they demonstrate that synonymous variants are much less likely than nonsynonymous variants to be deleterious. Overall, the report by Shen at al. does not provide sufficient grounds for overturning the large body of existing knowledge on the relative importance of synonymous and nonsynonymous variation.

## Extended Data

**Extended Data Figure 1.**
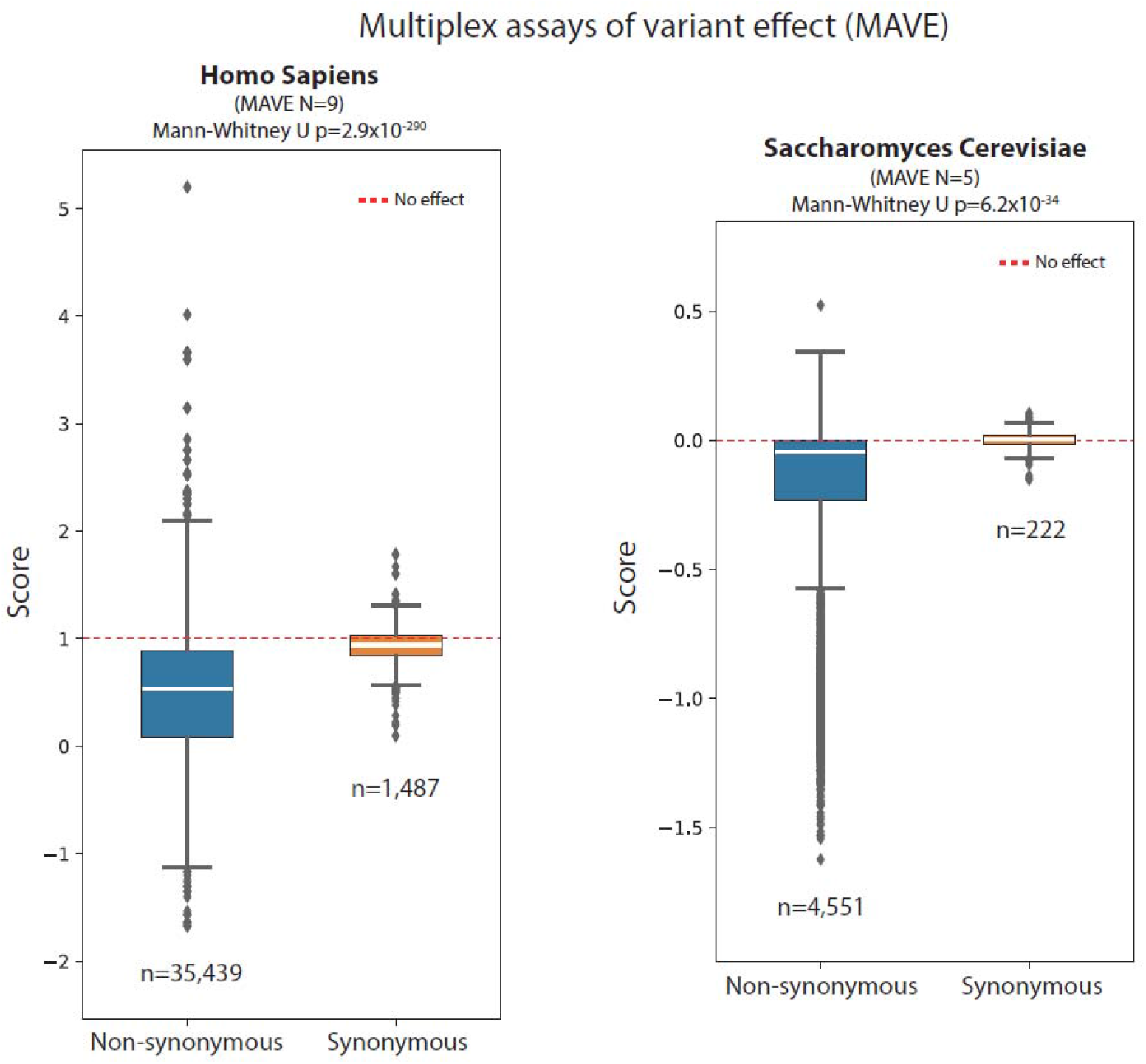
The effects of nonsynonymous and synonymous variants in multiplexed assays of variant effects. Abundance scores for MAVEs performed for 8 human genes (left) and 5 yeast genes (right) included in MaveDB^13^. For the human MAVEs, abundance scores were calculated based on a min-max normalization using wild type (score of 1) and the average nonsense variant score (score of 0). For yeast, effect sizes reflect the log2 ratio of each variant’s count to the wild type count.

## Methods

### UK Biobank

The UKB is a prospective study of approximately 500,000 participants aged 40–69 years at time of recruitment. Participants were recruited in the UK between 2006 and 2010 and are continuously followed. Participant data include health records that are periodically updated by the UKB, self-reported survey information, linkage to death and cancer registries, collection of urine and blood biomarkers, imaging data, accelerometer data and various other phenotypic end points. All study participants provided informed consent and the UK Biobank has approval from the North-West Multi-centre Research Ethics Committee (MREC; 11/NW/0382).

In this study, we analyzed exome and phenotypic data for 454,668 UK Biobank participants using our previously published pipeline^19^. Briefly, we excluded sequences that achieved a VerifyBAMID freemix (measure of DNA contamination) of more than 4% and samples where less than 94.5% of the consensus coding sequence (CCDS release 22) achieved a minimum of ten-fold read depth. We excluded participants that were second-degree relatives or closer.

In terms of phenotypic data, we analyzed the February 2020 data release that was subsequently refreshed with updated Hospital Episode Statistics (HES) and death registry data by the UKB in July 2020 (UKB application 26041). As previously described, we grouped relevant ICD-10 codes into clinically meaningful “Union” phenotypes.

### Site frequency spectrum analysis

We compared the allele frequency distributions of synonymous, missense, and protein-truncating variants (PTVs) observed in 454,668 UK Biobank participants in the same manner as our previously published analysis of the gnomAD dataset^11^.

On the basis of SnpEff annotations, we defined synonymous variants as those labeled as synonymous_variant and restricted to only fourfold degenerate codons. We defined missense variants as those labeled missense_variant. We defined PTVs as variants annotated as exon_loss_variant, frameshift_variant, start_lost, stop_gained, stop_lost, splice_acceptor_variant, splice_donor_variant, gene_fusion, bidirectional_gene_fusion, rare_amino_acid_variant, and transcript_ablation.

For synonymous variants, we also annotated codon optimality changes using Codon Stability Coefficient (CSC) scores derived from HEK293T cells^12^. Consistent with our prior analysis, we classified synonymous variants that resulted in changes from a codon with a positive CSC to a negative CSC as “optimal-to-nonoptimal” (O > NO), the opposite as “nonoptimal-to-optimal” (NO > O), and all others as “neutral.” We compared differences in allele frequency distributions using a two-tailed Fisher’s Exact Test.

### Phenome-wide association study

We retrieved gene-phenotype association statistics from our publicly accessible portal of gene-based collapsing analyses performed on UK Biobank exomes (https://azphewas.com)^19^. These collapsing analyses were performed on a subset of 394,692 UK Biobank participants of European ancestry. The carriers of at least one qualifying variant (QV) in a gene were compared to the non-carriers using a two-tailed Fisher’s exact test for 10 different collapsing models that capture a range of genetic architectures. Here, we specifically focused on three different qualifying variant collapsing models: the synonymous model, the “raredmg” missense model, and the PTV model. In the synonymous model, qualifying variants (QVs) were defined as synonymous variants with a gnomAD MAF <= 0.005%, UKB MAF <= 0.05%. QVs in the raredmg model included missense variants with gnomAD MAF <= 0.005%, UKB MAF <= 0.1%, and a REVEL score >= 0.25. The PTV model included variants with gnomAD MAF <= 0.1% and UKB MAF <= 0.1%. The average number of QVs per individual was 35.17 for the synonymous model, 28.4 for the “raredmg” model, and 8.6 for the PTV model. Thus, the synonymous model was equally, if not more, powered for the detection of gene-phenotype associations.

### Compilation of published MAVE datasets for Homo sapiens and Saccharomyces cerevisiae

We employed the MaveDB^13^ public repository to retrieve datasets from Multiplexed Assays of Variant Effect (MAVEs) both for Homo Sapiens and Saccharomyces cerevisiae. We first downloaded all available score sets for each species via the MaveDB API and filtered out any datasets that did not provide informative scores for synonymous variant effects (i.e. provided a fixed value of 1, other constant value, or no value for all synonymous variants as a representative default null model for any further analysis). In order for the results to be comparable, we retained for Homo Sapiens those score sets that were based on min-max normalization using wild type (score of 1) and the average nonsense variant score (score of 0), leading to 9 datasets in total (accession numbers: urn:mavedb:00000001-a-3, urn:mavedb:00000001-a-4, urn:mavedb:00000001-b-2, urn:mavedb:00000001-c-1, urn:mavedb:00000001-d-1, urn:mavedb:00000013-a-1, urn:mavedb:00000013-b-1, urn:mavedb:00000095-a-1, urn:mavedb:00000095-b-1), covering 8 genes (*UBE2I, SUMO1, TPK1, CALM1, CALM2, CALM3, PTEN*, and *CYP2C9*).

For Saccharomyces cerevisiae, we retained all score sets reporting effect sizes as log2 ratio of each variant’s count to the wild type count, leading to 5 datasets overall (accession numbers: urn_mavedb_00000011-a-1_scores.csv, urn_mavedb_00000037-a-1_scores.csv, urn_mavedb_00000039-a-1_scores.csv, urn_mavedb_00000040-a-1_scores.csv, urn_mavedb_00000074-a-1_scores.csv). For each dataset, the “hgvs_pro” was used to infer the effect type of each variant (i.e., synonymous or missense); only synonymous and missense variants were included in our analysis. The “score” field from each dataset was used to compare the distribution of effects from synonymous vs missense variants. The statistical significance of the difference between the distributions was quantified with Mann-Whitney U test.

## Data availability

Association statistics generated in this study are publicly available through our AstraZeneca Centre for Genomics Research (CGR) PheWAS Portal (http://azphewas.com). All whole-exome sequencing data described in this paper are publicly available to registered researchers through the UKB data access protocol. Exomes can be found in the UKB showcase portal: https://biobank.ndph.ox.ac.uk/showcase/label.cgi?id=170. MAVE data was accessed through mavedb.org.

## Acknowledgements

We thank the participants and investigators in the UKB study who made this work possible (Resource Application Number 26041). We thank Dr. Craig Kaplan and Dr. Chirag Vasavda for helpful comments on the manuscript.

